# Ovicidal Activity of 2-Hydroxy-4-Methoxybenzaldehyde, Derivatives and Structural Analogues on *Anopheles gambiae* eggs

**DOI:** 10.1101/2021.09.14.460396

**Authors:** R. E Andati, M. O. Omolo, I.O. Ndiege

## Abstract

**Background:** Effective remedies for disrupting *Anopheles gambiae* metamorphosis at the egg stage are crucial in suppression of the malaria vector populations that result in the reduction of disease burden. 2-Hydroxy-4-methoxybenzaldehyde (the major component of *Mondia whytei* roots), its derivatives, structural analogues and their blends were evaluated against the eggs of *An. gambiae* in the search for ovicidal compounds with potential use in mosquito control programs.

**Methods:** Mature roots were harvested from *Mondia whytei* plants grown in the Center for African Medicinal & Nutritional Flora and Fauna (CAMNFF) herbal medicinal garden and cleaned with distilled water. 2-Hydroxy-4-methoxybenzaldehyde (**1**) was isolated by steam distillation of the chopped roots. The selected derivatives and/or analogues were prepared using established chemical procedures and their structures confirmed by NMR spectroscopy and ESI-MS. Ovicidal activity of the pure compounds, derivatives, structural analogues and/or formulated blends was tested at 1, 10, 25 and 50 ppm on *An. gambiae* eggs..

**Results:** Eleven mono-substituted (**3-7**), di-substituted (**8-10**), tri-substituted (**1-2**) aromatic compounds were assayed for ovicidal activity against *Anopheles gambiae* eggs singly or as blends. Benzaldehyde (**4**) and 4-methoxybenzaldehyde (**9**) were further converted into 2-hydroxy-1, 2-diphenylethanone (**11**), 1, 5-diphenylpenta-1, 4-diene-3-one (**12**) and 1, 5-*bis* (4-methoxyphenyl) penta-1, 4-diene-3-one (**13**) and evaluated for ovicidal activity individually or as blends. Of the thirteen compounds evaluated individually, 2-hydroxy-4-methoxybenzaldehyde (**1**) exhibited the highest ovicidal activity at LC_50_ 0.7075 ppm while anisole had the lowest activity at LC_50_ 40.342 ppm. The derivatives exhibited moderate activity: 2-hydroxy-1, 2-diphenylethanone (LC_50_ 10.599 ppm), 1, 5-diphenylpenta-1, 4-diene-3-one (LC_50_ 9.019 ppm) and 1, 5-bis (4-methoxyphenyl) penta-1, 4-diene-3-one (LC_50_ 15.642 ppm). The blends exhibited intriguingly high ovicidal efficacy with the mixture of benzaldehyde and phenol showing the highest (LC_50_ 0.332 ppm) while phenol and anisole exhibited the lowest activity (LC_50_ 9.9909 ppm).

**Conclusion:** From the activity of the blends, it is evident that anisole is antagonistic to the efficacy of phenol and benzaldehyde. It is also apparent that aldehyde and hydroxyl groups, when directly attached to the phenyl ring, provide the critical structural characteristics that contribute to the ovicidal activity of the aromatic compounds.

## Introduction

Malaria remains the most important parasitic disease in the world [1]. Africa with an estimated 215 million annual malaria cases accounts for 94% of the global cases leading to 384,000 deaths [2]. It is estimated that there were 33 million pregnancies in Africa in 2020 with 35% of the expectant mothers being exposed to malaria infection resulting in about 82,000 children with low birth weight [2].

Mosquitoes are important public health vectors of malaria, filariasis and arboviral diseases that cause millions of infections and death worldwide [3]. Malaria is transmitted by infected female *An. gambiae* which feed on human blood meal for viability of its eggs [4]. Effective vector control methods at the egg, larval or adult stages are therefore critical in controlling the malaria vector and mitigating its harmful effects on human health [5]. Most malaria control strategies: environmental management (breeding/resting sites), sterile insect technique: biological control agents (predators, parasitoids and entomopathogens); chemical repellents and insecticide/pesticides (natural and synthetic), depend heavily on insect vector population control of the larval or adult stages with little effort on the eggs [6]..

Natural insecticide/pesticides are generally non-pest specific, biodegradable, non-allergic to humans, safe to non-target organisms [7] and have wide spectrum of application [8]. They offer good alternatives to synthetic chemical insecticides which are deleterious to the environment, harmful to non-target organisms and are ineffective due to development of resistance [9]-[13]. Many reports of plant extracts, secondary metabolites, essential oils and lectins which exhibit: general insect toxicity; growth and/or reproductive inhibition; insect repellency; and larvicidal activity against mosquito vectors have been documented; and are important and potentially suitable for use in integrated vector management (IVM) [6], [14].. However, little work has been documented on ovicidal activity and oviposition deterrence of anti-mosquito plants and/or derived compounds.

In that regard, the ovicidal activity and oviposition deterrency of leaf extracts of *Ipomoea cairica* against dengue vectors [6]; ovicidal and repellent activity of several botanical extracts against *Culex quinquefasciatus, Aedes aegypti* and *Anopheles stephensi* [14]; and the ovicidal and larvicidal activity of some plant extracts against *Cx. quinquefasciatus* and *Ae. aegypti* [13] have been documented. The larvicidal, ovicidal and oviposition deterrent potential of neem oil water dispersible tablets have been reported against *An. culicifacies* [15] while azadirachtin from neem plant exhibited ovicidal activity against *Cx. tarsalis* and *Cx. quinquefasciatus* [16].. The larvicidal and ovicidal activity of *Artocarpus blancoi* extracts were noted against *Ae. aegypti* [17] while the larvicidal, ovicidal, and repellent activity of *Sophora alopecuroides* and synergistic activity of its dominant constituents have been documented against *Ae. albopictus* [18].

Efforts to control malaria transmission in disease endemic areas are heavily reliant on suppression of the vector populations through a combination of chemicals, biological methods and management of breeding sites [19]. Consequently, application of adulticides and larvicides has been a common strategy used in vector control. It is of essence that focus be also directed to the egg stage in the mosquito development cycle due to its limited movement compared to the free flying and swimming adult mosquitoes and larvae, respectively [20]. Consequently, discovery and development of effective and environmental-friendly ovicidal compounds alongside the identification and focus on most productive/viable breeding sites/ habitats for mosquito is crucial for malaria vector and disease control [21].

2-Hydroxy-4-methoxybenzaldehyde (**1**), a structural isomer of vanillin (**2**), is an aromatic taste-modifying compound commonly found in the root bark of *Mondia Whytei* plant [22]. It has been previously reported as a tyrosine inhibitor [23] and potent larvicide against *An. gambiae* [24]. However, we could not find any information on its ovicidal activity or any structure-activity relationship studies on it or related compounds against *An. gambiae* eggs in the literature. Consequently, this project was designed to investigate the ovicidal activity of 2-hydroxy-4-methoxybenzaldehyde, its derivatives, structural analogues and their blends on *An. gambiae* eggs in order to understand the structure-ovicidal activity relationships therein.

## Materials and methods

### Insect culture

The *An. gambiae* mosquitoes that produced eggs used in this study were reared under ambient conditions of 27±1 °C and 85% relative humidity (RH), in the insectary situated at Centre for Disease Control CDC at the Kenya Medical Research Institute, Kisumu, Kenya. They were fed on 10% sucrose solution and a blood meal, to ensure that they produced viable eggs for the ovicidal experiments.

### Compounds for ovicidal assay

2-Hydroxy-4-methoxybenzaldehyde (**1**) was isolated from *Mondia whytei* Skeels while compounds **2-10** were procured from LOBA CHEMIE PVT LTD.

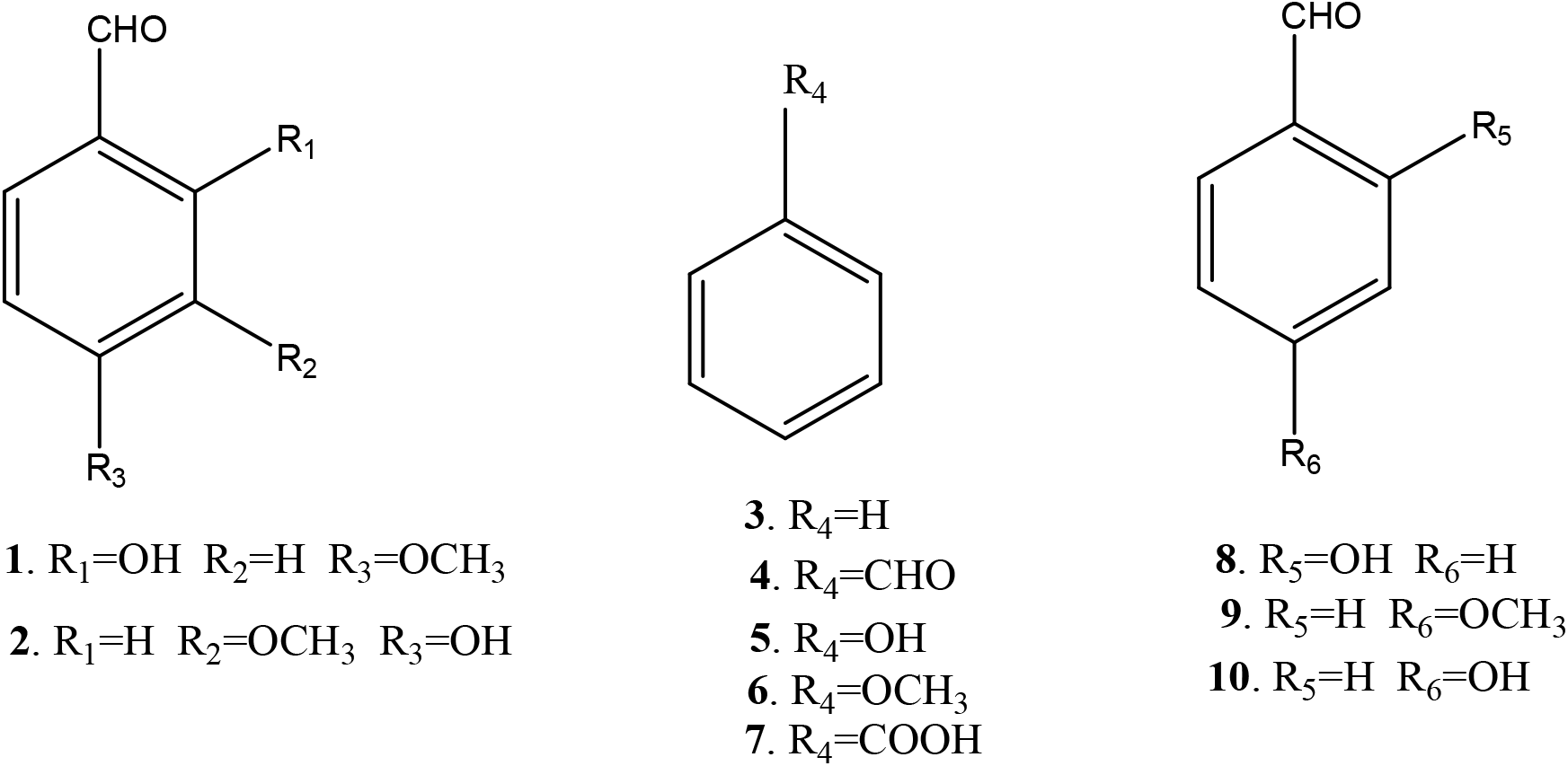

Compounds **11-13** were prepared in the laboratory as described below.

### Isolation of 2-hydroxy-4-methoxybenzaldehyde

Fully developed roots were harvested from mature *Mondia whytei* plants cultivated at the Centre for African Medicinal & Nutritional Fauna & Flora (CAMNFF) herbal garden at Masinde Muliro University of Science & Technology, Kakamega, Kenya. They were cleaned with water and stored under shade awaiting extraction. The roots were chopped into small pieces and subjected to isolation using the established procedures [25]. The white crystalline compound [10 g, mp 41-43 °C] was obtained from 1000 g (10% yield) of the roots and confirmed to be 2-hydroxy-4-methoxybenzaldehyde (**1**) from NMR and ESI-MS data [22]. It was stored in sealed amber bottles and refrigerated at 4 °C awaiting ovicidal assays.

### 2-Hydroxy-1, 2-diphenylethanone (11)

The compound was prepared through established procedures summarized in Scheme 1 [26].

**Scheme 1:**
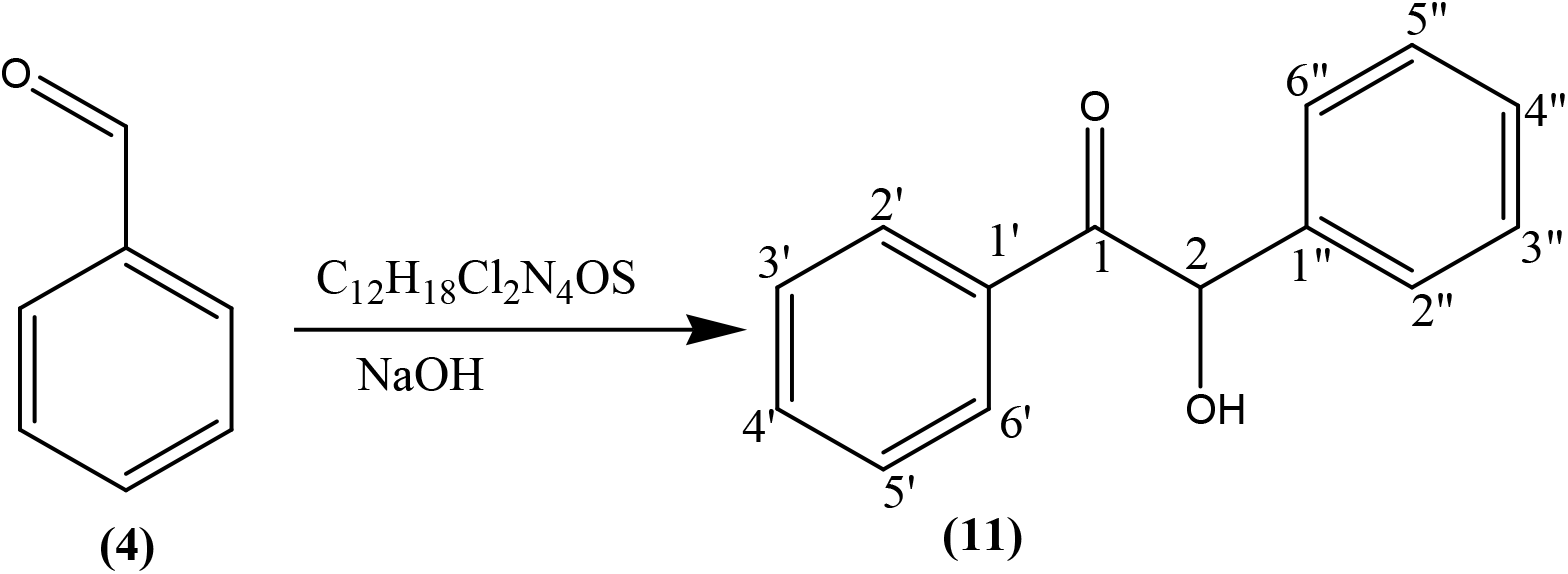
Preparation of 2-hydroxy-1, 2-diphenylethanone **(11)**

Briefly, thiamine hydrochloride (3 g, 8.04 mMol.) was dissolved in distilled water (4.5 mL) in a 250 mL conical flask, ethanol (30 mL) was added to the solution and the contents of the flask swirled by hand for 15 min until homogeneity was achieved. Addition of NaOH (9 mL, 0.25 Mol.) solution turned the mixture from colorless to bright yellow and the flask put on a mechanical shaker at 300 rpm for 20 min until the bright yellow color changed to pale yellow. Benzaldehyde (9 mL, 9.5 g, 85 mMol.) was slowly added to the mixture and the flask loaded onto a mechanical shaker at 300 rpm for 20 min until the mixture became homogeneous, the flask stoppered and left to stand in the dark for 72 hrs. The yellow crystals (8.25 g, 38.9 mMol.) were re-crystallized from hot ethanol to give white needle-like crystals of 2-hydroxy-1, 2-diphenylethanone (**11**) (7.92 g, 37.4 mMol., 95% yield) as confirmed by physical and spectroscopic data: mp 135-137 °C (literature 134-136 °C) [26]); ^1^H NMR (400 MHz, CD_3_SO) δ 7.82 (2H, d, J=5.16, H-2’,H-6’), 7.46 (2H, t, J=4.28, H-3’, H-5’), 7.76 (1H, m, H-4’), 7.39 (2H, d, J=6.8, H-3”, H-5”), 7.25 (2H, m, J=7.8, H-2”, H-6”) 7.06 (1H, d, J=5.08, H-4”); 5.92 (1H, d, H-2), 3.17 (1H, s, OH); ^1^H-^1^H COSY (CD_3_SO) see Figure 1; ^13^C NMR (δ, CD_3_SO) 199.65 (C-1) 140.2 (C-1”), 135.22 (C-1’), 133.69 C-4’), 128.93 (C-2’, C-6’), 128.17 (C-2’, C-6’) 129.3 (C-2”, C-6”), 129.06 (C-3”, C-5”), 127.73 (C-4”), 76.14 (C-2); DEPT 135 (CD_3_SO) 133.69 (C-4’H-4’), 128.17 (C-2” & C -6”H-2” & H-6”), 129.06 (C-3” & C-5”H-3” & 5”), 128.93 (C-2’ & C-6’H-2’ & H-6’), 127.73 (C-4”H-4”), 128.93 (C-2’ & C-6’H-2’ & 6’), and 76.14 (C-2H-2); ^1^J C-H, HSQC (CD_3_SO) C-2’ H-2’, C6’ H-6’,, C-3’ H-3’ C-5’H-5’ C-3” H-3” C-5”H-5”, C-4” H-4”, C-2” H-2” C-6”H-6”, C-4” H-4”, C-2 H-2; ^3^J, ^4^J H-C HMBC (CD_3_SO): see Figure 2; EIMS (*m/z*) 39, 51, 63, 77, 105 (100%) [C_7_H_5_O]^+^, 139, 165 210 [M^+^].

**Figure 1:**
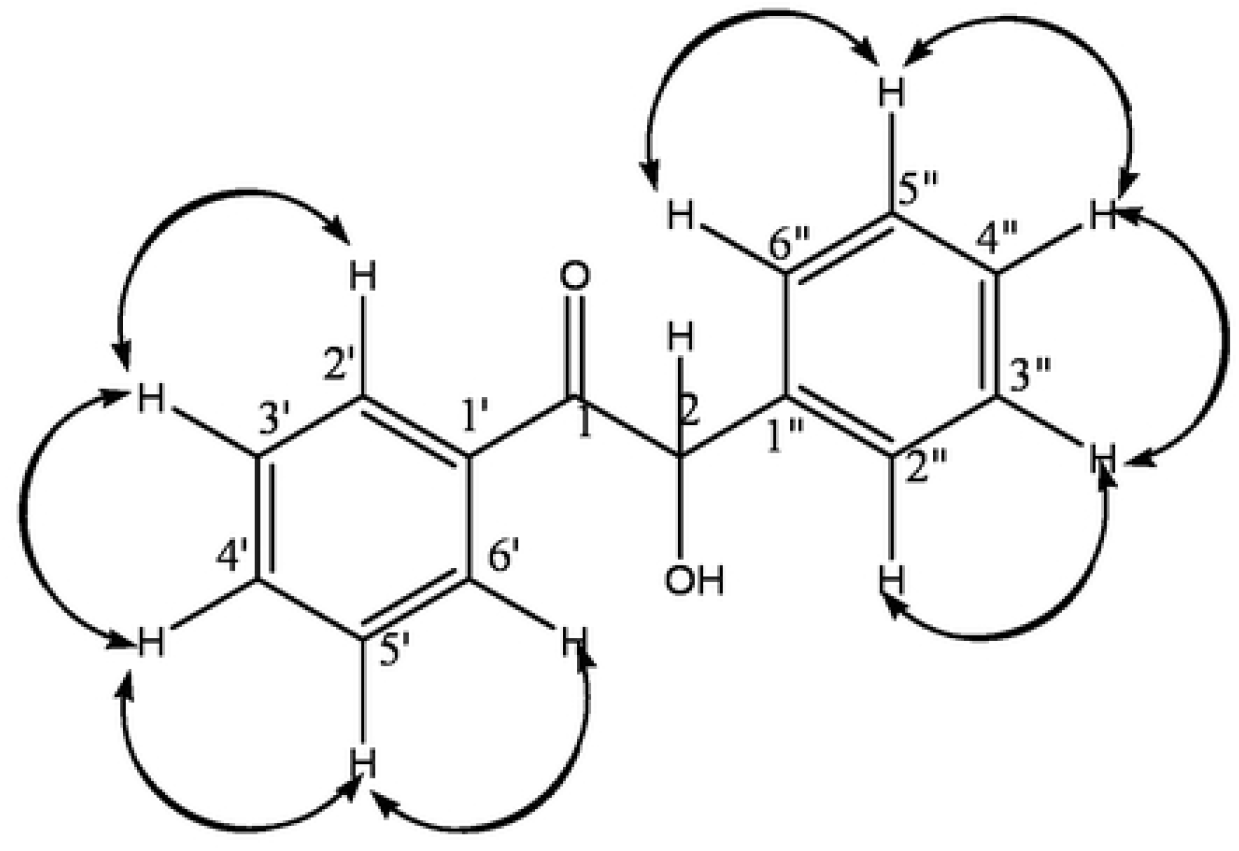
^1^H-^1^H COSY data for 2-Hydroxy-l, 2-diphenylethanone (**11**)

**Figure 2:**
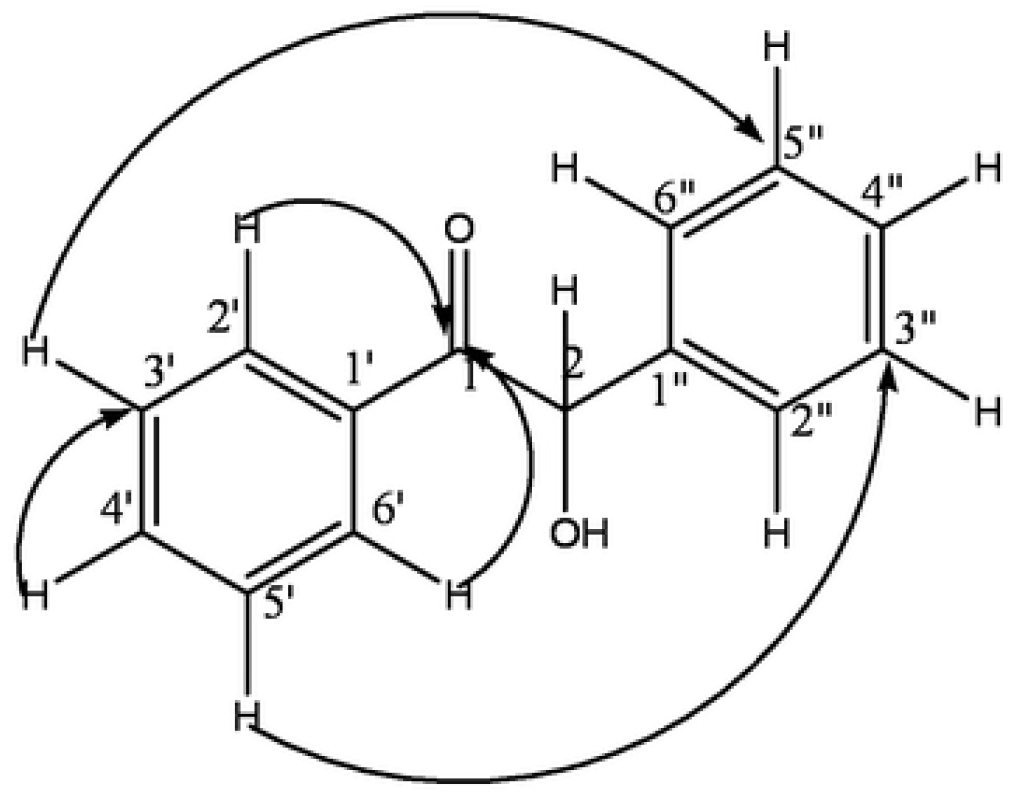
^3^J, ^4^J H-C HMBC data for 2-Hydroxy-1, 2-diphenylethanone (**11**)

Compounds **12** and **13** were prepared through the established procedures summarized in Scheme 2 [27].

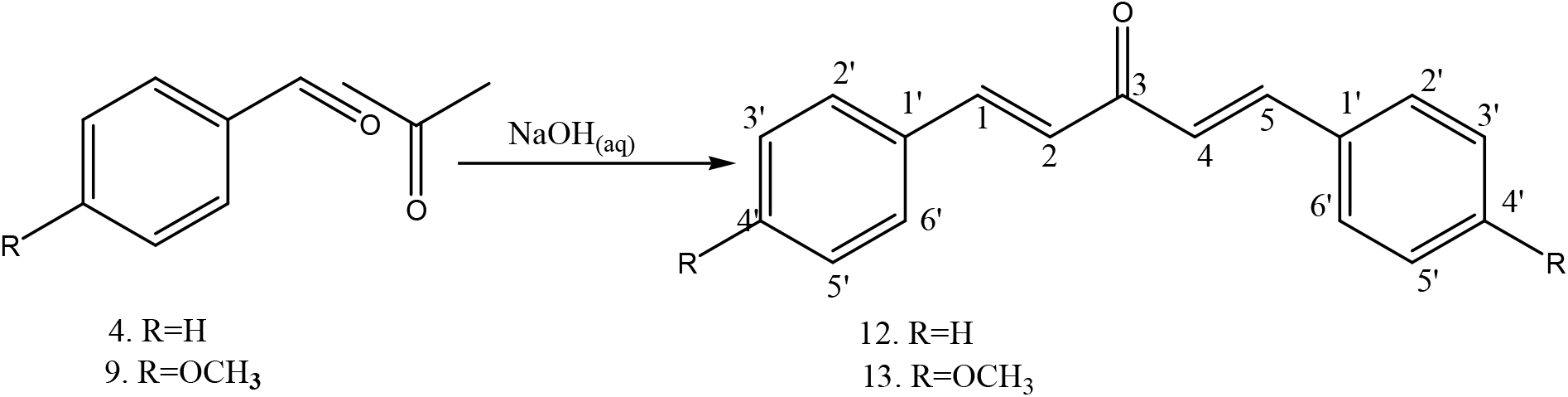

### 1, 5-Diphenylpenta-1, 4-diene-3-one (12]])

Briefly, 0.25 M NaOH solution (100 mL,) was transferred into a 500 mL conical flask, ethanol (80 mL) added and the mixture loaded onto a mechanical shaker at 200 rpm for 15 min. to attain homogeneity. Acetone (4 mL, 3.16 g, 54.4 mMol.) and benzaldehyde (12 mL, 12.47 g, 117.5 mMol.) were added and shaken at 200 rpm for 20 min while the mixture changed from pale yellow to deep yellow. On settling down, two layers were observed with the yellow crystals in the organic layer. The layers were separated, the organic layer filtered using a suction pump, the crystals collected, and dried to give 8.90 g. The crystals were carefully cleaned with methanol to obtain shiny disk like yellow crystals of 1.5-diphenylpenta-1,4-diene-3-one (**12**) (8.74 g, 37.5 mMol., 92% yield) and identified by physical and spectroscopic methods: mp 109-111 °C (literature 108-110 °C) [27]; ^1^H NMR (δ, CD_3_OD) 7.61 (4H, d, J=16.18, H-2’, H-6’), 7.32 (4H, d, J=8.04, H-3’, H-5’), 7.16 (2H, t, J=8.2, H-4’), 6.32 (2H, d, J=16.12, 12.64, H-2, H-4), 7.49 (2H, d, J=16.12, H-1, H-5); ^1^H-^1^H COSY (CD_3_OD): see Figure 3; ^13^C NMR (δ, CD_3_OD) 190.11 (C-3), 143.8 (C-1, C-5), 134.8 (C-1’), 130.38 (C-4’), 128.7 (C-2’, C-6’), 128.3 (C-3’, C-5’), 124.97 (C-2, C-4); DEPT 135 (δ) 143.8(C-1H-1,C-5 H-5) 130.38 (C-4’H-4’), 128.7 (C-2’H-2’,C-6’ H-6’), 128.3 (C-3’H-’), 124.97 (C-2 H-2, C-4 H-4); ^3^J, ^4^J H-C HMBC: see Figure 4; and EIMS (*m/z*) 39, 51, 63, 77, 91, 103, 115, 131, 156, 191, 205, 215,., 233 (100%) [M^+^-H], 234 [M^+^]

**Figure 3:**
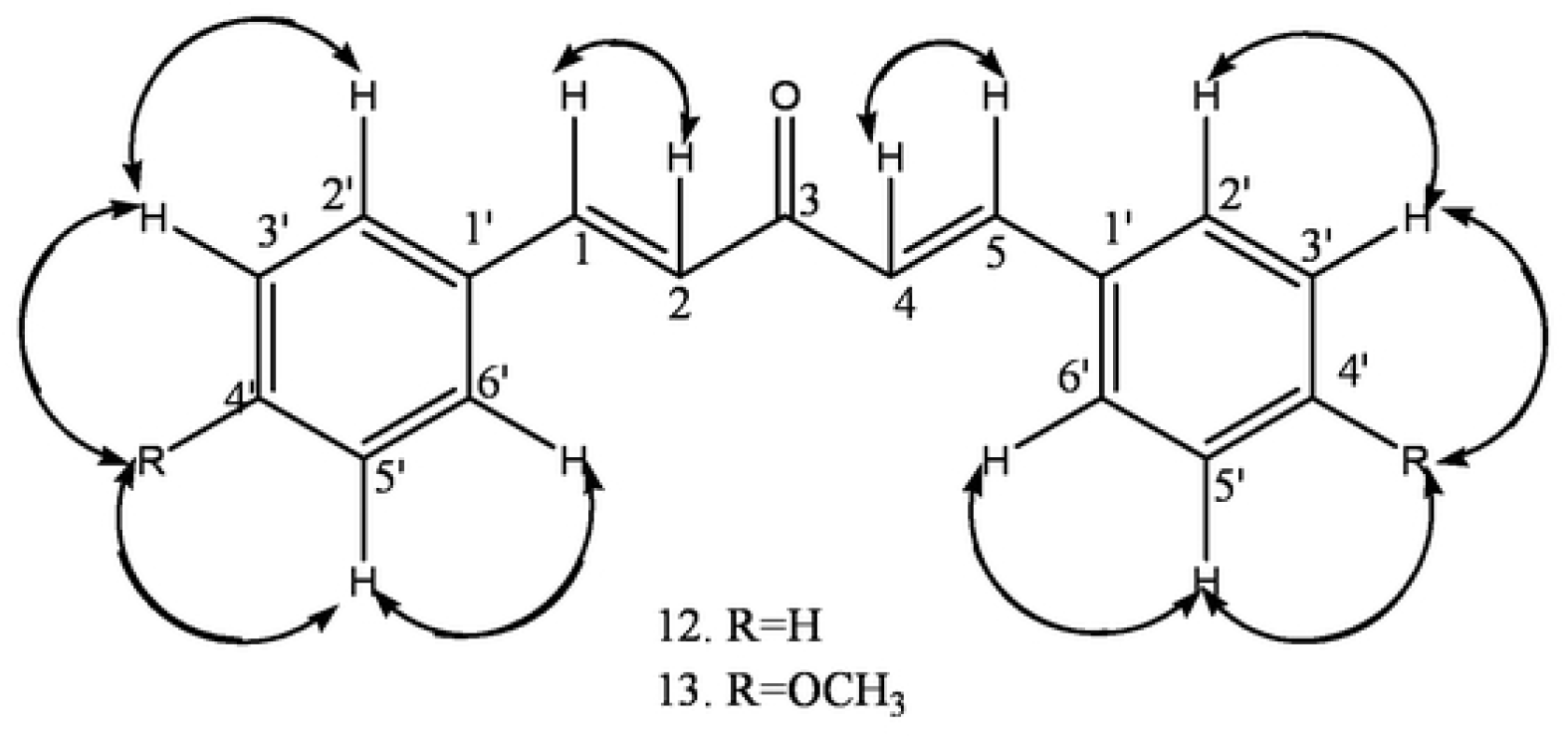
^1^H-^1^H COSY data for compound **12** and **13**

**Figure 4:**
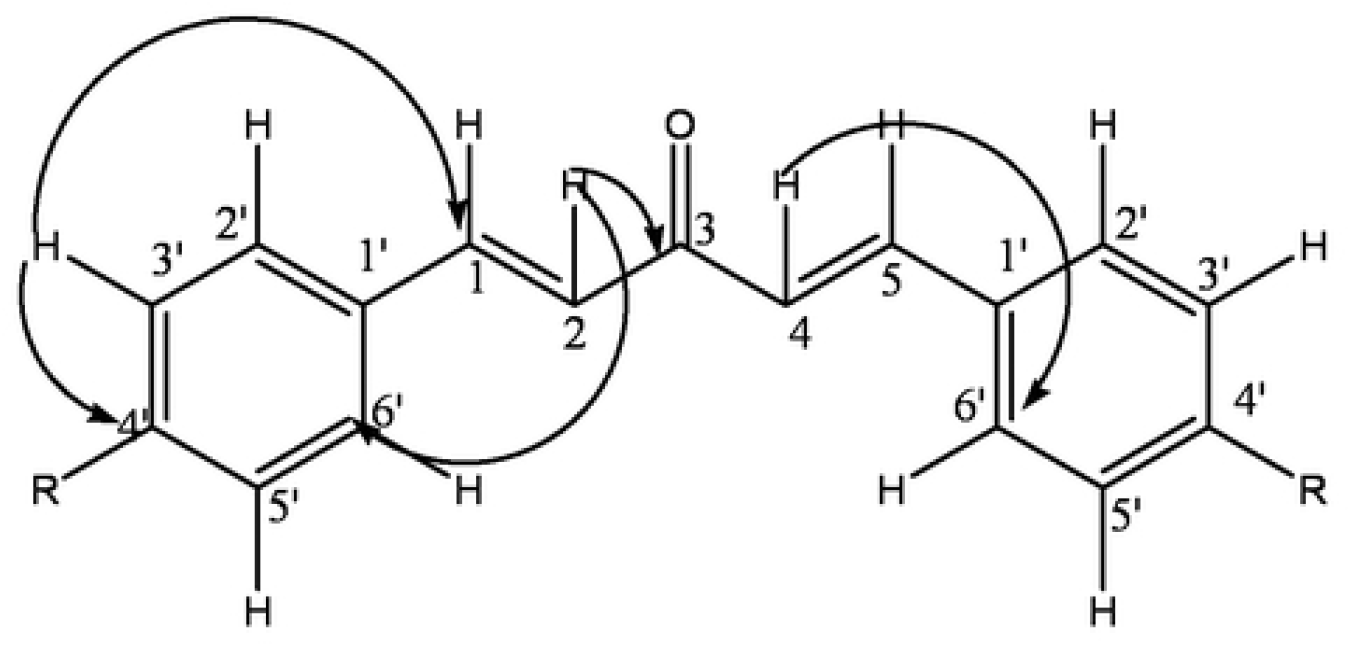
^3^J, ^4^J H-C HMBC data for compound **12 & 13**

### 1, 5-*Bis* (4-methoxyphenyl) penta-1, 4-diene-3-one (13)

Briefly, 0.25 M NaOH solution (25 mL) was poured into a 100 mL conical flask, 10 mL of ethanol added, and the mixture shaken at 200 rpm for 15 min. to attain homogeneity after which acetone (1.4 mL, 1.11 g, 19 mMol.) and 4-methoxybenzaldehyde (5.3 mL, 5.51 g, 40 mMol.) were sequentially added to form a white emulsion. The emulsion was shaken at 200 rpm for 10 min., allowed to settle down and form two layers: pale yellow and deep yellow, with fat-like droplets on top of the deep yellow layer, which changed into yellow crystals after 30 min. The mixture was filtered using a suction filtration pump to afford yellow crystals which were dried and cleaned carefully with methanol to give shiny disk-like yellow crystals of 1.5-*bis*-(4-methoxyphenyl)penta-1,4-diene-3-one **(13)** (3.83g, 13 mMol, 85% yield): mp 105-107 °C (literature 105-107 °C) [28]; ^1^H NMR (δ CDCl_3_) 7.89 (4H, d, J=8.8, H-2’, H-5’,), 7.65 (4H, d, J=8.8, H-3’, H-5’,), 7.42 (2H, d, J=15.8, H-1, H-5), 6.99 (2H, d, J=15.8, H-2 H-4). 3.87 (3H, s, OCH_3_); ^1^H-^1^H COSY (CDCl_3_): see Figure 5; ^13^C NMR (δ, CDCl_3_) 188.86 C-3) 161.57 (C-4’), 142.67 (C-1, C-5), 130.07 (C-2’, C-5’), 127.68 (C-1’), 123.54 (C-2, C-4), 114.43 (C-3’, C-5’), 55.41 (OCH_3_); DEPT 135 (δ, CDCl_3_) 142.67(C-1H-5), 130.07 (C-2’H-2’), 130.07 (C-5’, H-5’), 123.54 (C-2H-2), 114.43 (C-3’,H-3’), 114.43 (C-5’H-5’), 55.41 (OCH_3_); ^1^J C-H, HSQC (δ, CDCl_3_) C-2 H-2 4C-4 H-4, C-1 H-1 C-5H-5, C-3’ H-3’, C-5’H-5’, C-2’ H-2’; ^3^J, ^4^J H-C HMBC: see Figure 6; EIMS (*m/z*)[31, 39, 41, 53, 56, 57, 59 (100%) [C_3_H_4_O]^+^, 63, 73, 83, 87, 100, 294 [M^+^].

### Preparation of stock solution and dilutions

The pure compounds (10 mg) were dissolved in 10 mL of ethanol and topped up to 100 mL with distilled water to prepare 100 ppm stock solutions. The stock solutions were diluted appropriately with distilled water to obtain 1, 10, 25 and 50 ppm solutions for ovicidal assays.

Blends were formulated from benzaldehyde (**4**) (B), phenol (**5**) (P) and anisole (**6**) (A). Briefly, the individual compounds (3.33 mg each) were mixed to form blend BPA, which was dissolved in ethanol (10 mL) and topped up to 100 mL of distilled water to make a stock solution of 100 ppm. For blends PA, BP and BA, equal amounts of individual compounds (5 mg) were mixed and dissolved in 10 mL of ethanol and topped up to 100 mL with distilled water to make 100 ppm stock solutions. The stock solutions were diluted appropriately with distilled water to make 1, 2, 10, 25 and 50 ppm for ovicidal assays.

### Ovicidal activity

Ovicidal activity was determined by measuring the inhibition of egg hatchability. Briefly, freshly laid eggs of *An. gambiae* were counted and divided into groups of 100 using a hand magnifying lens and each group submerged into 25 mL of 1, 2, 10, 25 and 50 ppm solutions of each pure compounds in transparent plastic containers for 48 hours or until they hatched into larvae or completely inhibited from hatching. Each treatment was replicated 4 times, with eggs exposed to 1% ethanol in water and plain distilled water serving as controls. The hatchability was assessed after 48 hours of post treatment. The emergent larvae were also observed for survival rate and deformities.

### Statistical analysis

The ovicidal data were compared using Students Newman Kuel *t*-tests (SNK t-test) and the dose-response-relationships determined using probit analysis. The LD_50_ and LD_90_ values obtained from the regression analysis [29]. The level of significance of statistical data was set at *p* < 0.05 or lower.

## Results and discussion

Thirteen compounds (**1-13**) grouped into five molecular structures (**1-2, 3-7, 8-10, 11, 12-13**) were tested individually and as blends for ovicidal activity using the eggs of *Anopheles gambiae* mosquitoes.

The results in table 1 indicate the hatchability rate of the *An. gambiae* eggs in different treatments at various concentrations. Highest egg mortality or un-hatchability was observed at 50 ppm in nearly all the compounds tested. Poor ovicidal activity was noted for all the treatments at 1 ppm since almost all of the *An. gambiae* eggs hatched into larvae. The larvae that emerged had no significant deformities observed.

**Table 1:** Means (Mean ± SD) Number of Hatched Eggs of *An. gambiae* from Treatment with Compounds and Blends on at Various Concentrations.

| Compound/Blend | Concentration |  |  |  |
| --- | --- | --- | --- | --- |
|  | 1 ppm | 10 ppm | 25 ppm | 50 ppm |
| **1** | 44.33±5.93<sup>c</sup> | 0.0±0.0<sup>h</sup> | 0.0±0.0<sup>h</sup> | 0.0±0.0<sup>h</sup> |
| **2** | 75.00±8.90<sup>a</sup> | 83.25±7.05<sup>a</sup> | 68.75±4.89<sup>a</sup> | 0.0±0.0<sup>h</sup> |
| **3** | 67.50±1.85<sup>a</sup> | 83.25±7.05<sup>a</sup> | 48.25±5.54<sup>c</sup> | 0.0±0.0<sup>h</sup> |
| **4** | 59.00±3.74<sup>b</sup> | 69.75±3.15<sup>a</sup> | 0.0±0.0<sup>h</sup> | 0.0±0.0<sup>h</sup> |
| **5** | 84.25±3.86<sup>a</sup> | 45.00±3.35<sup>a</sup> | 2.00±1.23<sup>g</sup> | 0.0±0.0<sup>h</sup> |
| **6** | 92.00±2.38<sup>a</sup> | 54.00±3.54<sup>a</sup> | 49.25±1.93<sup>c</sup> | 0.0±0.0<sup>h</sup> |
| **7** | 80.25±8.35<sup>a</sup> | 66.50±5.27<sup>a</sup> | 76.00±6.34<sup>a</sup> | 0.0±0.0<sup>h</sup> |
| **8** | 69.75±4.50<sup>a</sup> | 0.0±0.0<sup>h</sup> | 0.0±0.0<sup>h</sup> | 0.0±0.0<sup>h</sup> |
| **9** | 62.67±17.84<sup>a</sup> | 0.0±0.0<sup>h</sup> | 0.0±0.0<sup>h</sup> | 0.0±0.0<sup>h</sup> |
| **10** | 85.75±3.50<sup>a</sup> | 75.00±2.58<sup>a</sup> | 19.50±3.88<sup>e</sup> | 0.0±0.0<sup>h</sup> |
| **11** | 67.00±7.94<sup>a</sup> | 57.50±3.71<sup>b</sup> | 19.00±3.49<sup>e</sup> | 0.75±0.75<sup>g</sup> |
| **12** | 76.00±9.44<sup>a</sup> | 55.75±8.43<sup>b</sup> | 0.75±0.48<sup>g</sup> | 0.0±0.0<sup>h</sup> |
| **13** | 85.00±3.50<sup>a</sup> | 75.00±2.58<sup>a</sup> | 19.00±3.88<sup>e</sup> | 0.0±0.0<sup>h</sup> |
| **BPA** | 57.67±16.19<sup>a</sup> | 25.00±11.00 | 4.67±3.71<sup>g</sup> | 0.0±0.0 |
| **BA** | 75.33±6.74<sup>a</sup> | 69.00±3.605<sup>a</sup> | 0.0±0.0 | 0.0±0.0 |
| **BP** | 45.67±4.91<sup>c</sup> | 16.33±10.398<sup>e e</sup> | 0.0±0.0 | 0.0±0.0 |
| **PA** | 56.33±23.82<sup>a</sup> | 41.67±3.180<sup>d</sup> | 14.33±2.03<sup>f</sup> | 0.0±0.0 |
| Ethanol | 71.25± 4.96<sup>a</sup> | 73.00±1.958<sup>a</sup> | 78.75±3.01<sup>a</sup> | 57.50±6.86<sup>b</sup> |
| water | 83.00±4.55<sup>a</sup> |  |  |  |

Means with the same letter are not significantly different (SNK test, p=0.0001).

### Structure activity relationship

2-Hydroxy-4-methoxybenzaldehyde **(1)**, a trisubstituted aromatic compound with hydroxyl, methoxyl and aldehydic groups, has been previously isolated from the roots of *Mondia whytei*, shown to be as a tyrosine inhibitor [23] and a potential larvicide for *An. gambiae* [24]. The compound is also responsible for the characteristic smell and taste of roots from the plant [22]. It is the reported insecticidal properties and the diverse funtional groups that prompted us to probe its ovicidal activity against *An. gambiae* eggs to establish whether it has potential to inhibit the eggs from hatching and if so which functional groups confer the activity. The ovicidal activity of 2-hydroxy-4-methoxybenzaldehyde (**1**), related compounds (**2-10**), structural analogues (**11-13**) and formulated blends against *An. gambiae* are summarized in table 2. With an LD_50_ value of 0.7075 ppm, compound **1** prompted a structure activity relationship to probe the functional groups responsible for the observed ovicidal efficacy. Readily avaible and a closely related congener; 4-hydroxy-3-methoxybenzaldehyde/ vanillin (**2**) was bioassayed and the activity found to be 34 times lower (LD_50_ 24.177 ppm) than **1**). Interestingly, lower tyrosinase inhibition and larvicidal activity of vanillin **(2)** and other related congeners have also been reported [23-24]. The big difference in the biological activity of the two compounds is quite intriguing given that they have similar functional groups save for their relative positions to each other on the aromatic skeleton. This observation prompted further investigations on the cause of varied activity in regard to the observed ovicidal activity. Consequently, related compounds with similar functional groups but different functional group arrangement on the benzene skeleton were assayed for comparison.

**Table 2:**
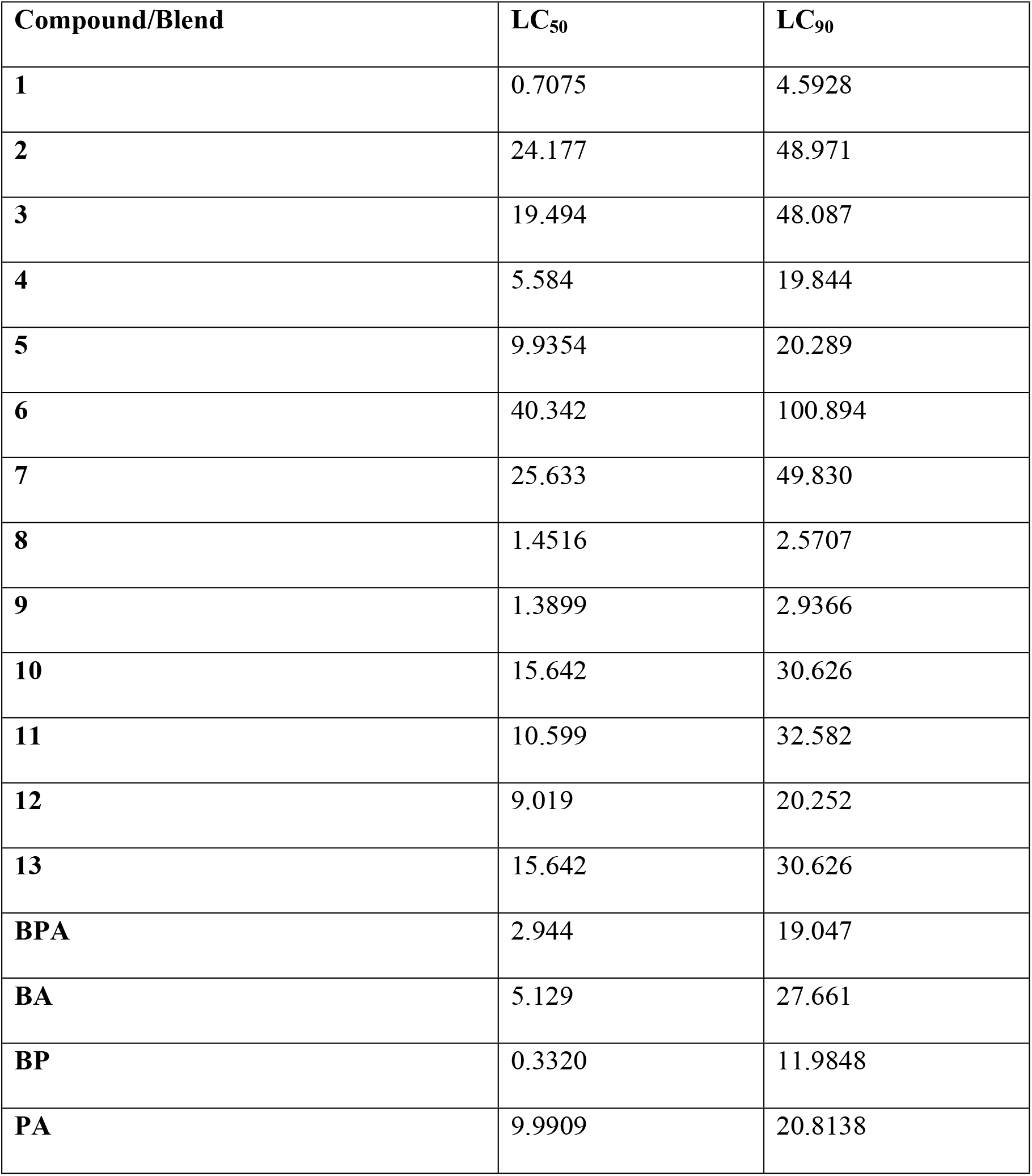
Ovicidal Activity (LC_50_ and LC_90_) of Evaluated Compounds and Blends on *An. gambiae* eggs.

Simple aromatic compounds constituting similar functional groups like those on **1** and **2** were assayed. Anisole or methoxybenzene (**6)** (LD_50_ 40.342 ppm), benzoic acid **(7)** (LD_50_ 25.633 ppm) and benzene (**3**) (LD_50_ 19.494 ppm) exhibited low activity while benzaldehyde (**4)** at LD_50_ 5.584 ppm had slightly higher activity than phenol (**5)** (LD_50_ 9.9354 ppm). The results revealed an interesting trend in the potency of derivatives of benzene (**3**) due to substituent variation that enhance or lower activity of the resulting aromatic compound. Further, they demonstrate that the aldehyde functional group is more potent when attached to the benzene ring than carboxylic acid as in compound **7** that resulted in much lower ovicidal activity than **4**. This observation is consistent with earlier reports where 4-methoxysalicyclic acid was found to exhibit lower larvicidal activity than 2-hydroxy-4-methoxybenzaldehyde [24]. The enhanced ovicidal activity of compounds **4** and **5** suggest that attachment of an aldehyde or hydroxyl group on **3** enhances its efficacy. On the hand, methoxyl and carboxylic acid groups on benzene ring lowers the ovicidal activity of the resultant compound drastically. While the bioassay data of these simple compounds were supposed to help us understand the individual contribution of the individual functional groups on the activity of 2-hydroxy-4-methoxybenzaldehyde (**1**) and vanillin (**2**), they could not satisfactorily explain the observed activity of compounds **1** and **2**, and therefore more assays using di-substituted aromatic compounds were undertaken to investigate the structure-activity relationships in the two compounds. Due to relatively strong ovicidal activity, benzaldehyde (**4**) was chosen as the starting point for the structure-activity studies. The introduction of an electron donating hydroxyl group at *ortho* position in compound **4** resulted in 2-hydroxybenzaldehyde (**8**) with improved ovicidal activity (LD_50_ 1.452 ppm), confirming that *ortho*-hydroxyl group synergizes ovicidal activity of benzaldehyde. However, when the hydroxyl group was shifted to *para* position as in 4-hydroxybenzaldehyde (**10**), the activity was drastically lowered to LD_50_ 15.642 ppm, confirming that *para*-hydroxyl group is antagonistic to the ovicidal activity of benzaldehyde. The two observations unravel the contribution of the hydroxyl group on the activity of **1** and **2** and confirm that the relative position of the substituents on the benzene skeleton is critical. Interestingly, addition of a stronger electron donating methoxy group at the *para* position of **4** gives 4-methoxybenzaldehyde (**9**) with increased activity at LD_50_ of 1.390 ppm. The activity of compound **9** helped us to explain the low ovicidal activity observed in vanillin (**2)**, a tri-substituted aromatic compound. Considering the observed low activity of 4-hydroxybenzaldehyde (**10)**, addition of methoxy group *ortho* to the hydroxy and *meta* to the carbonyl gives vanillin (**2**) with lower activity. It is interesting to note that shifting the hydroxyl of 2-hydroxybenzaldehyde (**8**) to *para* lowers the activity of the resulting 4-hydroxybenzaldehyde (**10**), further addition of methoxyl *ortho* to the hydroxyl of 4-hydroxybenzaldehyde (**10**), gives vanillin (**2**) with much lower activity than 4-hydroxybenzaldehyde (**10**). The trend in the activity of vanillin (**2**), 2-hydroxybenzaldehyde (**8**) and 4-hydroxybenzaldehyde (**10**) assisted in assessing the effect of substituent position on the aromatic ring. It further demonstrates that the hydroxyl and methoxyl groups are either synergistic or antagonistic to the ovicidal effect of the carbonyl when on the same ring depending on the position of attachment. In addition, the hydroxyl and methoxyl groups are inactive when *ortho* relative to each other as earlier reported for larvicidal activity [24]. On the contrary, 2-hydroxy-4-methoxybenzaldehyde (**1**) displays quite an interesting trend in activity. Considering 2-hydroxybenzaldehyde (**8**) and 4-methoxybenzaldehyde (**9**), the addition of methoxy at *para* to the carbonyl and *meta* to the hydroxyl of **8** increases the activity of **1**. Similarly, the addition of hydroxyl *ortho* to the carbonyl and *meta* to the methoxyl group in 4-methoxybenzaldehyde (**9**) gives 2-hydroxy-4-methoxybenzaldehyde (**1**) with increased activity suggesting that methoxyl and hydroxyl groups are potentiating the benzaldehyde group depending on their positions relative to the carbonyl and to each other when all the three functional groups are on the same benzene ring

The electron donating hydroxyl and methoxyl groups gave interesting results when attached at *para* to aldehyde carbonyl. It is important to note the fundamental role played by the slightly bulky methoxyl in increasing activity as in 4-methoxybenzaldehyde (**9**); while on the other hand, the hydroxyl group in 4-hydroxybenzaldehye (**10**) lowered activity. The free *para*-hydroxyl group plays more antagonistic ovicidal role to the aldehydic carbonyl position than when it is at *ortho* position probably due to stronger inter-molecular H-bonding than the intra-molecular ones in **8** which enhance activity. These observations can also be rationalized by the stronger electron donating property of *para*-methoxy than *para* hydroxyl group and the intra-molecular hydrogen bonding to the carbonyl by the *ortho*-hydroxyl group. The enhanced activity of compounds **8, 9** and **10** therefore reflects the effectiveness of the position of hydroxyl and methoxyl groups in relation to aldehyde group as demonstrated in compounds **1** and **2** where it was noted that the position of methoxyl or hydroxyl has impact on activity.

Compound **4** was modified to **11** and **12** while compound **9** gave **13**. The structural analogs were assayed for ovicidal activity. Compounds **11 (**LD_50_ 10.599 ppm) and **12** (LD_50_ 9.019 ppm) exhibited lower activity exhibited lower ovicidal activity than the parent compound **4**. Similarly, compound **13** exhibited lower activity (LD_50_ 15.642ppm) than the parent compound **9**. The observations suggest that the aldehyde and hydroxyl groups are critical for ovicidal activity of *An. gambiae* eggs as previously reported for larvicidal activity [24].

The blends assayed for ovicidal activity included benzaldehyde (**4**), phenol (**5**), and anisole (**6**) (**BPA**); (benzaldehyde and anisole) (**BA**), (benzaldehyde and phenol) (**BP**) and (phenol and anisole) (**PA**). **BPA**, a blend of compounds **4**, **5** & **6** exhibited better ovicidal activity at LD_50_ 2.944 ppm than any of the individual components, but was four times lower than 2-hydroxy-4-methoxybenzaldehyde (**1)** and eight times higher than vanillin (**2**), indicating that the compounds exert synergy when in the blend. Blend **PA**, equivalent to subtraction of compound **4** from **BPA**, lowered its activity by almost half to LD_50_ 5.129 ppm thus indicating that benzaldehyde is a critical component of the blend. Blend **BP**, equivalent to substituting compound **6** with **4** or subtracting **6** from blend **BPA**, exhibited the highest ovicidal activity LD_50_ 0.332 ppm which was nine (9) times higher than that of blend **BPA** and fifteen (15) times higher than that of blend **PA** and confirmed that benzaldehyde is a critical component of the blend. Blend **BA**, equivalent to the subtraction of compound **5** from **BPA** or substitution of compound **5** from **BP**, exhibited the lowest ovicidal activity of all the blends at LD_50_ 9.990 ppm confirming that anisole is an antagonistic component of the blend. The observed results revealed that the synergistic interaction of the individual compounds is much stronger when the compounds are blended than when all the functional groups are incorporated in one compound suggesting that intra-molecular interactions have higher positive impact on ovicidal activity than the inter-molecular interactions.

Several structure-larvicidal activity relationships have been documented with all the studies linking functional groups of different compounds to the resulting activity of the compounds. For instance, acetyl derivatives of monoterpenoid compounds were reported to have high activity against the larvae of *Ae. aegypti* [30]. In another case, presence of hetero-atoms in the basic monoterpene structure for instance neoisopulegol reduced the potency of the compound. It was further noted that conjugated or *exo*-carbon-carbon double bonds and epoxidation increased larvicidal activity [31]. The larvicidal assay of eugenol and its derivatives revealed that the derivatives had lower activity [32]. Furthermore, it has been reported that conversion of phenol to diphenyl ether increased the activity against *An. gambiae* larvae [24].

## Conclusion

Finally our work established that the hydroxyl, methoxyl and aldehyde functional groups on an aromatic skeleton confer ovicidal activity when appropriately located in one compound but are strongly synergistic when in different molecules. 2-Hydroxy-4-methoxybenzaldehyde exhibited the highest ovicidal activity against *An. gambiae*, while anisole exhibited the lowest efficacy. Simple mono-functional compounds: benzaldehyde, phenol and anisole exhibited relatively low activity than when evaluated individually than when formulated as blends. Among the blends, blend **BP** exhibited the highest activity, while **BA** had the lowest efficacy. The presence of aldehyde and hydroxyl groups on mono-substituted benzene confers strong ovicidal activity while methoxyl group lowers activity. For di-substituted simple aromatic compounds, methoxyl group is an activity-potentiating group at *para* position to the aldehyde group and hydroxy when in ortho-position to the aldehyde.

## Acknowledgements

This work was supported by Masinde Muliro University of Science and Technology (MMUST) University Research Fund (URF). We appreciate Mr. Richard Amito and his team at the Kenya Medical Research Institute (KEMRI) for the supply of *An. gambiae* eggs and the help accorded to us during the bioassays. Center for African Medicinal and Nutritional Flora and Fauna (CAMNFF) is acknowledged for provision of *Mondia whitei* roots and the laboratory space for synthetic work.

## Captions

S1 Table 1: Means (Mean ± SD) Number of Hatched Eggs of *An. gambiae* from Treatment with Compounds and Blends on at Various Concentrations

S2 Table 2: Ovicidal Activity (LC_50_ and LC_90_) of Evaluated Compounds and Blends on *An. gambiae* eggs

S3 Figure 1: ^1^H-^1^H COSY data for 2-Hydroxy-1, 2-diphenylethanone (**11**)

S4 Figure 2: ^3^J, ^4^J H-C HMBC data for 2-Hydroxy-1, 2-diphenylethanone (**11**)

S5 Figure 3: ^1^H-^1^H COSY data for compound **12** and **13**

Figure 3: ^1^H-^1^H COSY data for compound **12** and **13**

S6 Figure 4: ^3^J, ^4^J H-C HMBC data for compound **12 & 13**

## Notes

### Competing Interest Statement

The authors have declared no competing interest.

## References

1. WHO (2010). World Malaria Report, Geneva: World Health Organization.

2. WHO (2020). World Malaria Report: 20 Years of Global Progress and Challenges: World Health Organization, Geneva; Licence: CC BY-NC-SA 3.0 IGO.

3. Senthil N. (2009). The use of *Eucalyptus tereticornis* Sm.(Myrtaceae) oil (leaf extract) as a natural larvicidal agent against the malaria vector *Anopheles stephensi* Liston (Diptera: Culicidae). Bioresource Technology 98: 1856–1860.

4. Takken W, Knols BGJ (1999). Odour mediated behaviour of afro-tropical malaria mosquitoes. Ann. Rev. Entomol. 44: 131–57.

5. WHO (1995). Vector Control for Malaria and Other Mosquito-borne Diseases. Report of a WHO Study Group. WHO Technical Report Series 857, World Health Organization, Geneva, Switzerland.

6. Ahbirami R., Zuharah W.F., Yahaya Z.S., Dieng H., Thiagaletchumi M, Fadzly N., et al. (2014) Oviposition deterring and oviciding potentials of *Ipomoea cairica* L. leaf extract against dengue vectors. Tropical Biomedicine 31: 456–465.

7. Bowers WS. (1992) Biorational approaches for insect control. Korean J. Appl. Entomol. 31: 289–303.

8. Ghosh .A, Chowdhury N., Chandra G. (2008). Laboratory evaluation of a phytosteroid compound of mature leaves of Day Jasmine (Solanaceae: Solanales) against larvae of *Culex quinquefasciatus* (Diptera: Culicidae) and non-target organisms. Parasitology Research. 103: 271–277.

9. Sutthanont N., Choochote W., Tuetun B.., Junkum A., Jitpakdi A., Chaithong U, et al. (2010). Chemical composition and larvicidal activity of edible plant-derived essential oils against the pyrethroid-susceptible and resistant strains of *Aedes aegypti* (Diptera: Culicidae). J. Vector Ecol. 35: 106–115.

10. Bayen S. (2012) Occurrence, bioavailability and toxic effects of trace metals and organic contaminants in mangrove ecosystems: a review. Environment International 48: 84–101.

11. Mulyatno KC, Yamanaka A, Ngadino, Konishi E. (2012). Resistance of *Aedes aegypti* (L.) larvae to temephos in Surabaya, Indonesia. S. E. Asian J. Trop. Med Pub. Health 43: 29–33.

12. Chavshin AR, Dabiri F, Vatandoost H, Bavani MM. (2015) Susceptibility of *Anopheles maculipennis* to different classes of insecticides in West Azarbaijan Province, Northwestern Iran, *Asian Pacif*. J. Trop. Biomed. 5: 403–6.

13. Munusamy R.., Appadurai D.R., Kuppusamy S., Michael G.P., Savarimuthu I. (2016). Ovicidal and larvicidal activities of some plant extracts against *Aedes aegypti* L. and *Culex quinquefasciatus* Say (Diptera: Culicidae). Asian Pacif. J. Trop. Disease 6: 468–471.

14. Govindarajan M., Mathivanan T., Elumalai K., Krishnappa K., Anandan A. (2011) Ovicidal and repellent activities of botanical extracts against Culex quinquefasciatus, *Aedes aegypti*, and *Anopheles stephensi*. (Diptera:Culicidae). *Asian Pac*. J. Trop. Biomed. 1: 43–48.

15. Kala S., Naika S.N., Patanjali P.K., Sogan N. (2019). Neem oil water dispersible tablet as effective larvicide, ovicide and oviposition deterrent against *Anopheles culicifacies*. S. Afr. J.Bot. 123: 387–392.

16. Su T., Mulla S. (1998) Ovicidal activity of neem product (azadirachtin) against *Culex tarsalis* and *Culex quinquefasciatus* (Diptera: Culicidae). J.Amer. Mosquito Control Assoc. 14: 204–209.

17. Pineda-Cortel M.R.B., Cabantog R.J.R., Caasi P.M., Ching C.A.D., Perez J.B.S., Godisan P.G.M., et al. (2019). Larvicidal and ovicidal activities of *Artocarpus blancoi* extracts against *Aedes aegypti*. Pharmaceutical Biology 57:, 120–124; DOI:10.1080/13880209.2018.1561727.

18. Shoukat R.F., Shakeel M., Rizvi S.A.H.,. Zhang J.Z.Y,. Freed S., X. Xu et al. (2020) Larvicidal, ovicidal, synergistic, and repellent activities of *Sophora alopecuroides* and its dominant constituents against *Aedes albopictus*. Insects 11: 246–259; DOI:10.3390/insects11040246.

19. WHO (2009) Dengue: Guidelines for Diagnosis, Treatment, Prevention and Control, New edition. World Health Organization, Geneva.

20. Elimam A.M., Elmalik K.H., Ali F.S. (2009). Larvicidal, adult emergence inhibition and oviposition deterrent effects of foliage extract from *Ricinus communis* L. against *Anopheles arabiensis* and *Culex quinquefasciatus* in Sudan. Tropical Biomedicine 26: 130–139.

21. WHO (1996) Report of the Consultation on key Issues in dengue vector control towards the operationalization of a global strategy. World Health Organisation, Geneva.

22. Mukonyi K.W., Ndiege I.O. (2001) 2-Hyrdoxy 4-methoxybenzaldehyde: aromatic taste modifying compound from *Mondia whitei* Skeels. Bull. Chem. Soc. Ethiopia 15:137–141.

23. Kubo I., Kinst-Hori I., (1999). 2-Hydroxy-4-methoxybenzaldehyde: a potent tyrosinase inhibitor from African medicinal plants. Planta Medica 65: 19–22.

24. Mahanga G.M., Akenga T.O., Lwande W., Ndiege I. O. (2005). 2-hydroxy-4-methoxybenzaldehyde: larvicidal structure activity studies. Bull. Chem. Soc. Ethiopia 61–68.

25. Rathi N., Keerthana H., Jayashree V., Sharma A. & Rao N.N. (2017) 2-Hydroxy-4-methoxybenzaldehyde, an astounding food flavoring metabolite: A review. Asian J. Pharmaceutic. Clin. Res.. 10: 105–110.

26. Rajeev S. M., Akkattu T. B., Vijay N. (2016). Recent advances in N-heterocyclic carbine (NHC)-catalyzed benzoin reactions. Beilstein J. Org. Chem. 12: 444–461, DOI: 10.3762/bjoc.12.47.

27. Nagaraja N., Vijay H. K, Swetha S. (2011). 1, 5-Diphenylpenta-1, 4-dien-3-ones: A novel class of free radical scavengers. Bulgarian Chem. Comm. 43: 460–464.

28. Harrison W. T. A., Sarojini B. K., Vijaya R. K. K., Yathirajan H. S., Narayana B. (2006). A redetermination of 1,5-*bis*(4-methoxyphenyl)penta-1,4-dien-3-one at 120 (2) K. Acta Cryst., E62, [1522–1523]; DOI: 10.1107/S1600536806009640

29. SAS (2012) SAS Guide for Personal Computers, Version 8.2. SAS University Edition.

30. Pandey S. K, Tandon S., Ahmad A., Singh A. K., Tripathi A.K. et al. (2013). Structure-activity relationships of monoterpenes and acetyl derivatives against *Aedes aegypti* (*Diptera: Culicidae*) larvae. Pest Manage. Sci. 69: 1235–123; DOI:10.1002/ps.3488.

31. Sandra R.L., Melo M.A., Valenca A., Santos R.L.C., De Sousa D.P., Socates C.H. (2011) Structure-Activity relationship of larvicidal monoterpenes and derivatives against *Aedes aegypti* Linn. Chemosphere 84:150–153; DOI: 10.1016/j.chemosphere.2011.02.018.

32. Barbosa J.D.F., Silva V., Alves P., Giuseppe G., Roseli L.C., Damiao P. et al. (2012). Structure-activity relationships of eugenol derivatives against *Aedes aegypti*. Pest manage Science 68(11):1478–83; DOI: 10.1002/ps.3331.

